# High predation risk decimates survival during the reproduction season

**DOI:** 10.1101/2022.03.31.486539

**Authors:** Radovan Smolinský, Zuzana Hiadlovská, Štěpán Maršala, Pavel Škrabánek, Michal Škrobánek, Natália Martínková

## Abstract

1. Predators attack conspicuous prey phenotypes that are present in the environment. Male display behaviour of conspicuous nuptial colouration becomes risky in the presence of a predator, and adult males face higher predation risk. High predation risk in one sex will lead to low survival and sex ratio bias in adult cohorts, unless the increased predation risk is compensated by higher escape rate.
2. Here, we tested the hypothesis that sand lizards (*Lacerta agilis*) have sex-specific predation risk and escape rate. We expected the differences to manifest in changes in sex ratio with age, differences in frequency of tail autotomy, and in sex-specific survival rate.
3. We developed a statistical model to estimate predation risk and escape rate, combining the observed sex ratio and frequency of tail autotomy with likelihood-based survival rate. Using Bayesian framework, we estimated the model parameters. We projected the date of the tail autotomy events from growth rates derived from capturerecapture data measurements.
4. We found statistically stable sex ratio in age groups, equal frequency of tail regenerates between sexes, and similar survival rate. Predation risk is similar between sexes, and escape rate increases survival by about 5%. We found low survival rate and a low number of tail autotomy events in females during months when sand lizards mate and lay eggs, indicating high predator pressure throughout reproduction. Our data show that gravid females fail to escape predation.
5. The risks of reproduction season in an ectotherm are a convolution of morphological changes (conspicuous colouration in males, body allometry changes in gravid females), behaviour (nuptial displays), and environmental conditions which challenge lizard thermal performance. Performance of endotherm predators in cold spring months endangers gravid females more than displaying males in bright nuptial colouration.

## Introduction

Wild organisms experience abiotic and biotic interactions in the environment they inhabit. One crucial trade-off is navigating the predation risk that reduces fitness by contributing to mortality, but also increases resource expenditure through evading predation. Animals avoid detection by using camouflage (Stevens and Merilaita 2011), or by shifting their activity from thermal or time niche of the predator (Webb and Whiting 2005; Chen et al. 2021). The concept of finding the optimal survival strategy in a predator-prey interaction leads to a behavioral cascade where predators and prey interact (Angilletta Jr. 2009; Brown and Vincent 1992; Stuart-Fox et al. 2003; Kuo and Irschick 2016), creating a density- and frequency-dependent predator pressure on different prey phenotypes.

The prey influences the arms race in the predator-prey dynamic through mechanisms facilitating escape from the predator attack. One such mechanism is autotomy, and many species are able to shed and regrow a body part (Maginnis 2006; Bateman and Fleming 2009; Cooper Jr. et al. 2004; Baban et al. 2022). In extreme cases, animals can shed and subsequently regenerate their whole bodies (Mitoh and Yusa 2021), but appendage autotomy is more common. The animal under attack can break off its appendage in an attempt to escape the predator. Lizards shed tails, and the tail thrashes after autotomy, potentially distracting the predator, and allowing the prey to escape the immediate attack (Naidenov and Allen 2021). From a short-term perspective, the animal with a broken tail can run faster compared to an animal with an intact tail, but locomotion of animals with tail autotomy is negatively impacted when jumping (Fernández-Rodríguez and Branã 2020; Gillis et al. 2009) or climbing vertical surfaces (Fleming and Bateman 2012). On a long-term scale, tail autotomy imposes high energy and resource costs that negatively influence survival through the cost of injury, infection risk, lower ability to survive future predator attacks, and energy relocation to the regenerate (Wilson 1992; Bateman and Fleming 2009). Following appendage regrowth, survival of the post-autotomy animal improves (Lin et al. 2017).

In lizards, frequency of tail autotomy differs intraspecifically between populations and between sexes. The different strategies that fine-tune tail autotomy frequency balance resource availability with intra- and inter-specific interactions. Populations exposed to higher predator pressure show lower frequency of tail autotomy, especially when the predators are efficient in killing the prey (Itescu et al. 2017). However, the propensity for autotomy is critical in escaping the attack and further increases with increased food availability, intraspecific competition and animal boldness (Kuo and Irschick 2016; Talavera et al. 2021).

While intraspecific competition increases tail autotomy frequency in males and not in females (Itescu et al. 2017; Kuo and Irschick 2016; Talavera et al. 2021), males also tend to face higher predation risk (Stuart-Fox et al. 2003). The rationale explaining why lizard sexes might face different predation risk and perform with different escape rate follows sex differences in morphology and behaviour. Sexually dimorphic species show differences in performance in laboratory experiments (Lailvaux et al. 2003; Massetti et al. 2017; Simon et al. 2022), where males are generally bigger and faster. In nature, sexual differences in performance exhibit complex adjustments that result in a performance outcome similar in both sexes. Males and females might operate at different body temperature, select different microhabitats, or utilize different antipredatory behaviour (Lailvaux et al. 2003; Vanhooydonck and Van Damme 2003; Samia et al. 2016; Gomes et al. 2018).

The synergy of performance integration in sexes might become challenged during reproduction. Males display bright nuptial colouration as a signal to potential mates and competitors (Olsson 1994; Marshall and Stewens 2014; Badiane and Font 2021). While location of the nuptial colouration and its light spectrum reflectance evolves as a conjugation of roles in crypsis from predators and signalling to conspecifics, males tend to be exposed to higher predation risk compared to females (Danchin et al. 2008; Stuart-Fox et al. 2003; McQueen et al. 2017). In females, body allometry and weight increase associated with gravidity decrease speed and agility compared to males (Bauwens and Thoen 1981; Shine 2003; Husak 2006). Females can compensate the increasing load during gestation by exerting more power and increasing cadence in running (Scales and Butler 1997; Irschick et al. 2003; Zamora-Camacho et al. 2014).

When behavioural and physiological compensations of different biological realities of the sexes are insufficient to maintain similar survival under predator pressure, we can expect changes in demographic parameters. We hypothesize that sexual differences in predation risk influenced by the escape rate will affect sex ratio, frequency of tail autotomy and survival rate (Table 1). Specifically, we predict that

**Table 1:** Hypotheses of predation risk and benefit of tail autotomy to escape from predation based on sex. Green – change of sex ratio between young and adult animals, blue – frequency of tail regenerates, red – sex-specific survival Rate;↑ – increase in males,↓ – decrease in males,∼ – no difference.

| in males | Higher predation risk |  |  | Equal predation risk |  |  |
| --- | --- | --- | --- | --- | --- | --- |
|  | sex ratio | autotomy | survival | sex ratio | autotomy | survival |
| Higher escape rate | ∼ | ↑ | ∼ | ↑ | ↑ | ↑ |
| Equal escape rate | ↓ | ∼ | ↓ | ∼ | ∼ | ∼ |

1. if males face higher predation risk, but also benefit from higher escape rate, we expect to observe stable sex ratio in different age cohorts and similar survival rate between sexes, but males will have higher frequency of tail autotomy.
2. If the escape rate does not differ between sexes, but males suffer from higher predation risk, males will have lower survival rate, and consequently, the sex ratio will be skewed in favour of females in older cohorts.
3. If predation risk does not differ between sexes, the putative higher escape rate in males will result in males dominating sex ratio in older cohorts, having suffered from tail autotomy more frequently, but also having higher survival rate.
4. We expect no difference in the observed parameters in the case where the predation risk and escape rate do not differ between sexes (Table 1).

Here, we study the influence of predation risk and escape rate on population dynamics and survival of sand lizards (*Lacerta agilis*, Lacertidae, Reptilia). The sand lizard is a small (*<* 19 cm), sexually dimorphic lizard, distributed in the temperate Palearctic. Males are brightly coloured in spring months, and their colour changes with progressing season to more dull colours (Fig. 1). Males in nuptial colouration show hues of green and brown on lateral and sometimes dorsal sides, and females and juveniles hues of brown.

**Figure 1:**
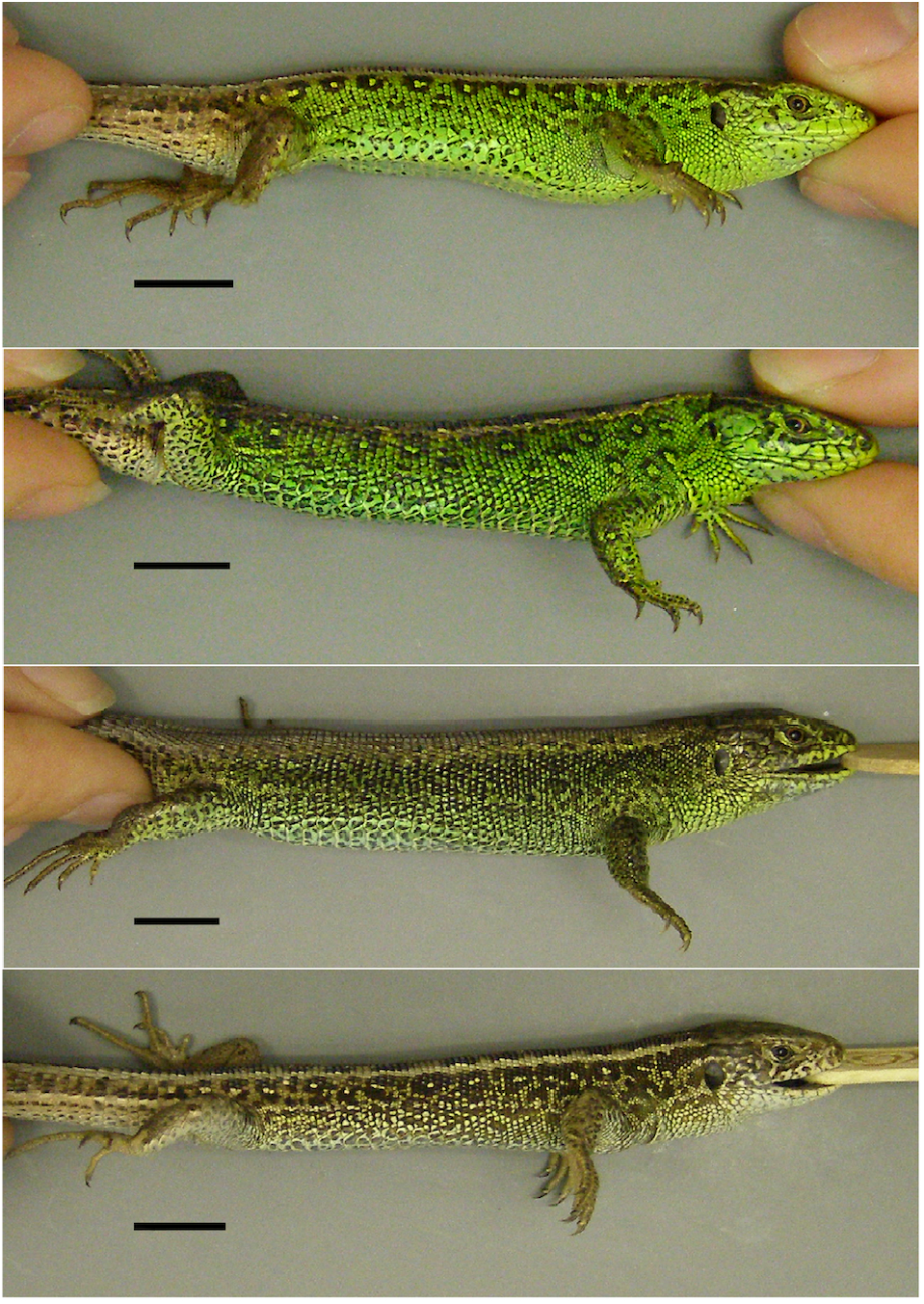
Sand lizard males (*Lacerta agilis*) show green nuptial colouration on their sides in spring (top panels). The colour changes to hues of brown with progressing season. Colour pattern is variable in the species, but seasonally changes only with respect to the size of the spots. Scale bar – 10 mm.

## Materials and methods

### Data

We sampled the sand lizard population during the months when the animals were apparent in an orchard in Hustopeče (48.93 N, 16.72 E) in Czechia. The orchard is on a southwest-facing slope, covers about 12 ha, and is surrounded by urban development and intensively managed fields. The area, where the sand lizards were sampled, is at the center of the orchard with dimensions 15 × 240 m and is mowed twice a year. Species considered sand lizard predators observed at the site include *Phassianus colchicus, Falco tinnunculus, Buteo buteo, Lanius collurio, Coronella austriaca, Mustela sp*. and domestic cats and dogs, with pheasants being frequently sighted. The sand lizards were apparent at the site between May and August 2018, May and September 2020 and 2021. We caught the animals by hand or by noosing (García-Muñoz and Sillero 2010) at weekly intervals with the mean number of days between trapping sessions equal to 8.2. We marked each captured individual by toe clipping or heat branding (Ekner et al. 2011). Prior to release, we photographed the animals on a photographic grey card with a ruler from dorsal, ventral and lateral sides (Fig. S1). We categorised the animals to age classes, sexed them, and collected data and material for other studies (Dračková et al. 2020; Smolinský et al. 2021). All sampling was performed by authorized personnel. Because the body size of individuals in age cohorts can vary (BÖhme and Bischoff 1984; Blanke and Fearnley 2015), we estimated the cohort according to size, colour and colour pattern described from Czechia (Gvoždík 2000; Smolinský et al. 2021). We considered the animals to be juveniles when they were captured between mid-summer and autumn, were small (body length from rostrum to cloaca (*L*_*ra*_) less than 47 mm), and had yellow ventral side with unresolved pattern on the dorsal side. The animals considered subadults were caught from spring to mid-summer, occasionally in late autumn, had *L*_*ra*_ *<* 66 mm, and started to develop colour pattern of adults. The remaining animals were considered adults. Juveniles and subadults were considered young animals in this study. At the site, sand lizards reach sexual maturity and enter reproduction after having overwintered twice (Gvoždík 2000; Smolinský et al. 2021).

We used the obtained capture-recapture data either by filtering the last known capture of each animal or by evaluating all captures of all individuals. Unless specified, the analyses use the non-redundant database of last captures of each animal.

### Age

We calculated the approximate age of sand lizards on the day of capture (*a*_*d*_ in days) according to

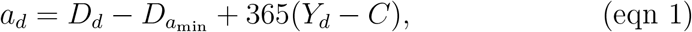

where *D*_*d*_ is the calendar day on the day of capture, *D*_*a*min_ is the calendar day of the first time a hatchling was encountered, *Y*_*d*_ is the year of capture, and the cohort *C* is the latest possible year, when the animal could have hatched. The cohort was equal to the year of first capture for juveniles, to the year previous to the year of first capture in subadults and to two years before the year of first capture in adults.

For example, let’s take a young and an adult animal captured for the first time on 1 June 2021. To estimate their ages, we need to know that the first hatchling was found on 1 August 2020 the previous year. The approximate age of the young sand lizard on the day of capture would be *a*_*d*_ = 152 − 214 + 365(2021 − 2020) = 303 days, because the young animal captured in spring had to have overwintered once and its year of hatching was 2020. The adult animal overwintered at least twice, bringing its year of hatching to, at most, 2019 and the corresponding approximate age to, at least, 668 days. To account for uncertainty in the day of birth of the individual animals, we recalculated the approximate age to the nearest month as *a*_*m*_ = 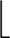*a*_*d*_*/*30.41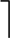, where 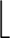•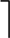 indicates rounding to the nearest integer and 30.41 is the average number of days in a calendar month. The animals in the example would thus be 10 and 22 months old, respectively.

### Sex

We sexed adult animals based on their nuptial colouration and presence of a swollen base of the tail. In young animals, we counted scales in the second rows from the ventral medial line (ventralia count) (Eplanova and Roitberg 2015). We developed a computer vision system to automate the counting of scales in digital photographs of sand lizard ventral sides. To establish a threshold for sexing juveniles, we counted the ventralia scales in adults and estimated the threshold value using the receiver operating characteristic curve (ROC), maximizing the specificity and sensitivity of detecting the correct sex. We validated the threshold values with logistic regression, and using the ROC analysis of ventralia count in young animals that were sexed upon recapture as adults.

Three centroid-based object detectors (Dolezel et al. 2022), which return the coordinates of scale centroids, form the core of the scale counting system (Fig. S1). The scale detectors utilize localization maps generated using UNet models (Ronneberger et al. 2015). Inputs and outputs of the models are 512 × 512 pixel (px) colour images and one-class localization maps, respectively. The target scales in the maps are represented as circles with centroids placed at coordinates corresponding to the centres of the scales. The circle circumferences represent the borders of nonzero values in the maps, where values at the centroid coordinates are at one, and the values within a circle decrease to zero with increasing distance from the circle centroid (Dolezel et al. 2022).

We trained and validated the U-Net models on a set of 227 expert-annotated photos, following the recommended training-validation procedure (Dolezel et al. 2022). The annotator marked a centre of each target scale in each photo. For each model, we generated a different set of localization maps using the centres (i.e. each model was trained on a different set of annotated photos). Specifically, we used maps with circles of 10 px diameters and linear decrease of gradients, maps with linear decrease of gradients and diameters respective to the size of scales (diameters 6–14 px), and maps with diameters respective to the size of scales but with nonlinear decrease of gradients.

To increase the accuracy of the scale detectors, we use square crops of uniformly oriented photos and maps as the inputs and outputs of the U-Net models, respectively. The crops are focused on the centres of lizard bodies. We utilize the YOLO object detector (Bochkovskiy et al. 2020) to create the crops (Fig. S1). The YOLO detector recognises and localizes heads, bodies and tails on the photographs. We use the acquired information about positions of the body parts to determine dimensions of the crops, and to rotate the photographs and the maps adequately. The size of each square crop is 1.42 times the length of the lizard body. To keep the aspect ratio of the crops, if necessary, the photos and the maps are padded with zeros before their resizing to 512 × 512 px. We trained and validated the YOLO detector on a set of 150 expert-annotated photos, following the recommended training-validation procedure (Bochkovskiy et al. 2020).

To count the scales at the left and the right side of the rows, we take into account subsequent positions of the centroids. We implemented all the processes (Fig. S1) into a custom algorithm ScalesCounter and used the algorithm to perform automated processing of unannotated photos. The algorithm is available upon request. Following the automatic counting, we checked all outputs and corrected the number of detected scales in low quality images (saturated image, appendage partially obscuring the scales, scarred scales).

### Seasonality

We considered two seasonal phases with different biological realities determining assumed activity and predation risk of sand lizards. Phase I started from the arousal from hibernation, and lasted until the first adult male started to lose nuptial colouration. Adult animals participate in reproduction in phase I, with males searching for mates and defending their territories, and females investing in gravidity. Phase II constituted the season after males started to lose nuptial colouration, and lasted until the beginning of hibernation. In phase II, the animals feed to accumulate sufficient resources for the winter.

### Tail autotomy

To estimate individual body and tail lengths, we used tpsDig v.2.02 (Rohlf 2005) software to obtain measurements from digital images of sand lizard ventral side. We measured body length from rostrum to cloaca (*L*_*ra*_), tail length from cloaca to tail tip (*L*_*cd*_), and length of the tail regenerate from break point to tail tip (*L*_*reg*_). We recorded a tail autotomy event, when the animal was first captured with a tail regenerate. Upon recapture, we compared the body measurements of the animals, and checked the photographic evidence for an additional tail autotomy event. If *L*_*cd*_ was shorter than on the day of the previous capture (with tolerance of 3 mm to account for muscle contraction), an additional tail autotomy event was recorded.

To calculate the date of the tail autotomy event, we first established individual recapture series starting with the record of the tail autotomy event and ending with the last capture in the same year as the first capture of the animal or before the additional tail autotomy event. We fitted linear models to the individual recapture series as *L*_*reg*_ = *β*_0_ + *β*_1_*a*_*d*_, where *a*_*d*_ is the approximate age of the animal (equation 1), *β*_1_ is the daily growth rate of the tail regenerate, and *β*_0_ is an intercept. We used median of individual tail regenerate growth rates at 10 mm intervals of *L*_*reg*_ to project the tail regenerate growth backward to estimate the approximate date of a tail autotomy event. The date was established for each recorded tail autotomy event independently.

## Statistical analysis

We tested the proposed hypotheses in two ways. First, we calculated the odds ratios for sex ratio in age groups and frequency of tail autotomy in sex and age groups, estimating the respective confidence intervals from likelihood profiles. For this, we used the last captures of each individual. Second, we calculated the survival rate and recapture probability in a likelihood framework based on the open population Cormack-Jolly-Seber (CJS) model (Cormack 1964; Jolly 1965; Seber 1965). For the survival rate analyses, we used the complete dataset that included the recaptures.

We calculated the odds ratio for the sex ratio change with age as

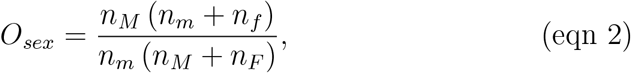

where *n* is the number of individuals, *M* and *F* represent males and females, respectively, with capital letters indicating adults and lower case letters indicating young animals.

We used data from adult and young animals separately to calculate the odds ratio for presence of tail regenerates. The change in odds of the presence of a tail regenerate in males compared to females is given as

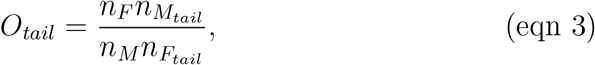

where 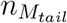 and 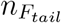are the numbers of males and females with tail autotomy, respectively.

To evaluate the change in the probability of a tail autotomy event with time, we modeled the relationship with logistic regression using sex, approximate age (in months) and their interaction as explanatory variables. The reported effect sizes are expressed as percent and given as 100(*O* − 1) for odds ratios and as 100(*e*^*β*1^ − 1) for logistic regressions, where *β*_1_ is the rate parameter.

We calculated the survival rate (*ϕ*) and recapture probability (*p*) from the capture-mark-recapture data, using the CJS model implemented in MARK (Cooch and White 2021). We used sex, age class (young, adult) and season (phase I, phase II) as categorical variables and tail length corrected for body length *L*_*cd*_*/L*_*ra*_ (mm mm^*−*1^) and *a*_*m*_ as individual covariates. All individual variable values were those recorded at first capture of each animal. We compared the models with AICc and corrected the parameter standard errors for overdispersion. We performed a goodness-of-fit test for the CJS model and tested for overdispersion with the median *ĉ* approach. The global model for the overdispersion test was *ϕ*(sex*age*season)*p*(sex*age*season), where season indicates the two seasonal phases.

### Predation risk and escape rate model

Sex differences in predation risk and escape rate will manifest in differences in survival. Survival rate *ϕ*, the probability that an individual will survive a specific time period, is the residual of mortality rate *m, ϕ* = 1 − *m*. The mortality rate can be attributed to predation (*m*_*P*_) and other causes (*m*_0_), giving *ϕ* = 1 − (*m*_0_ + *m*_*P*_). To reduce dimensionality of the problem, we set *m*_0_ = 0, and consider *ϕ* = 1 − *m*_*P*_. Mortality rate attributable to predation is predation risk (*P*) reduced by the escape rate (*E*), giving *m*_*P*_ = *P* − *E*. Escape rate can be thought of as a probability that increases survival, where the action of prey results in its immediate survival following the predator attack. While the prey can escape an attack unharmed, here we consider the escape to represent a situation in which the prey lost its tail. A predator attack that did not result in prey mortality or tail autotomy will not manifest in the parameters measured in this study. Substituting *m*_*P*_, the survival rate becomes

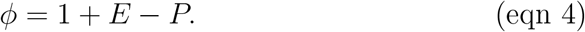

Sex-specific survival rate will influence how the proportion of each sex changes in time. The ratio of male (*ϕ*_*M*_) and female (*ϕ*_*F*_) survival rate will be equal to the odds ratio of sex ratio between age classes

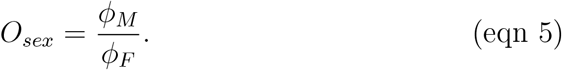

Substituting *ϕ* with equation 4, we get

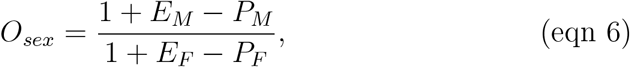

where the indices *M* and *F* indicate males and females, respectively.

The ratio of male-to-female escape rate should reflect the odds of frequency of tail autotomy in each sex. The relationship between sex-specific escape rate and odds ratio of tail autotomy in adults will be

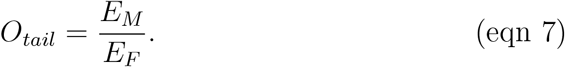

Estimating sex-specific predation risks and escape rates requires a four-dimensional optimisation for which an analytical solution might not exit. To solve the problem, we co-estimated the *P*_*M*_, *E*_*M*_, *P*_*F*_, and *E*_*F*_ using equations 4, 6 and 7 with a Bayesian approach. We calculated log-likelihood (ln *L*) of the observed data given the sex-specific predation risk and escape rate as

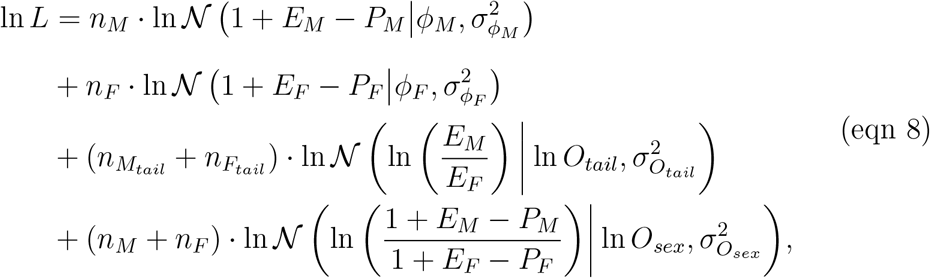

where 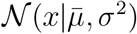 is the density of *x* in a normal distribution with mean 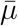 and variance *σ*^2^. We sampled the posterior with Metropolis-coupled Markov chain Monte Carlo (MCMC) using uninformative, uniform priors for sexspecific predation risks (*U* (0.001, 0.999)) and escape rates (*U* (0.001, 1.000)). We ran the MCMC for 5 mil. steps, sampling every 1000th to avoid autocorrelation, and discarding 20% of initial steps as burnin. We evaluated MCMC convergence from two independent runs with random starting values and combined the results.

We performed the statistical analyses in R (R Core Team 2020) using packages imager (Barthelme 2020), pROC (Robin et al. 2011), RMark (Laake 2013), plot3D (Soetaert 2019), coda (Plummer et al. 2006), and RColorBrewer (Neuwirth 2014).

### Ethics statement

Sampling was based on permits JMK 38000/2018 and JMK 42819/2021 issued by the Regional Authority of the South Moravian Region, Brno. Animal handling complied with Czech Act No. 114/1992 on Nature and Landscape Protection. The authors were authorised to handle wild lizards according to the Certificate of Professional Competence (Nos. CZ01287 and CZ03799; §15d, Act No. 246/1992 Coll.).

## Results

To estimate the approximate age of the sand lizards, we used dates when the first hatchlings of the season were seen. These were on 28 July 2018, 16 August 2020, and 29 July 2021. To estimate the seasonal phases, we found the seasonal phases breakpoints on 23 June 2018, 25 July 2020, and 2 July 2021, when the first male started to lose the green, nuptial colouration.

For sex and presence of tail autotomy to evaluate escape rate and predation risk in sand lizards, we investigated 365 individuals in total. To estimate the sex differences in ventralia count, we first scored the adults. The ventralia scales could be counted from photographs of 126 adult females and 105 adult males. In adult sand lizards, the ventralia count differed between sexes (Welch two-sample *t*-test: *t* = 19.7, *df* = 322.0, *p <* 0.001) with females having on average (± SD) 61.32 ± 2.18 ventralia scales, and males having 56.70 ± 2.17 ventralia scales (Fig. S2A). The area under the ROC was equal to 0.92 (Fig. S2B). At the threshold value of 58.5, established from the Youden’s index, the specificity of determining sex based on the ventralia count was equal to 0.85, the respective sensitivity being 0.86. The threshold value was supported by the logistic regression (*p* = 0.501; Fig. S2C). In young animals, in which the sex was determined when they were recaptured as adults, sex determination from ventralia count had specificity equal to 0.82 and sensitivity to 0.80 at the threshold value (Fig. S2B). Based on these results, we consider the ventralia count a sufficiently reliable sex-determination marker of sand lizard population from Hustopeče, and we used the threshold value of 58.5 to sex the juveniles.

In 22 individuals, we could not determine sex or the body measurements due to the poor quality of pictures, caused by auto-focus malfunction. The study includes 343 animals with complete data that were captured 506 times (Table 2, Fig. 2A).

**Table 2:** Number of individual sand lizards, *Lacerta agilis*. Animals with tail autotomy in parentheses.

|  | Sex | Young | Adults |
| --- | --- | --- | --- |
|  | Males | 48 (8) | 112 (68) |
|  | Females | 48 (16) | 135 (79) |

**Figure 2:**
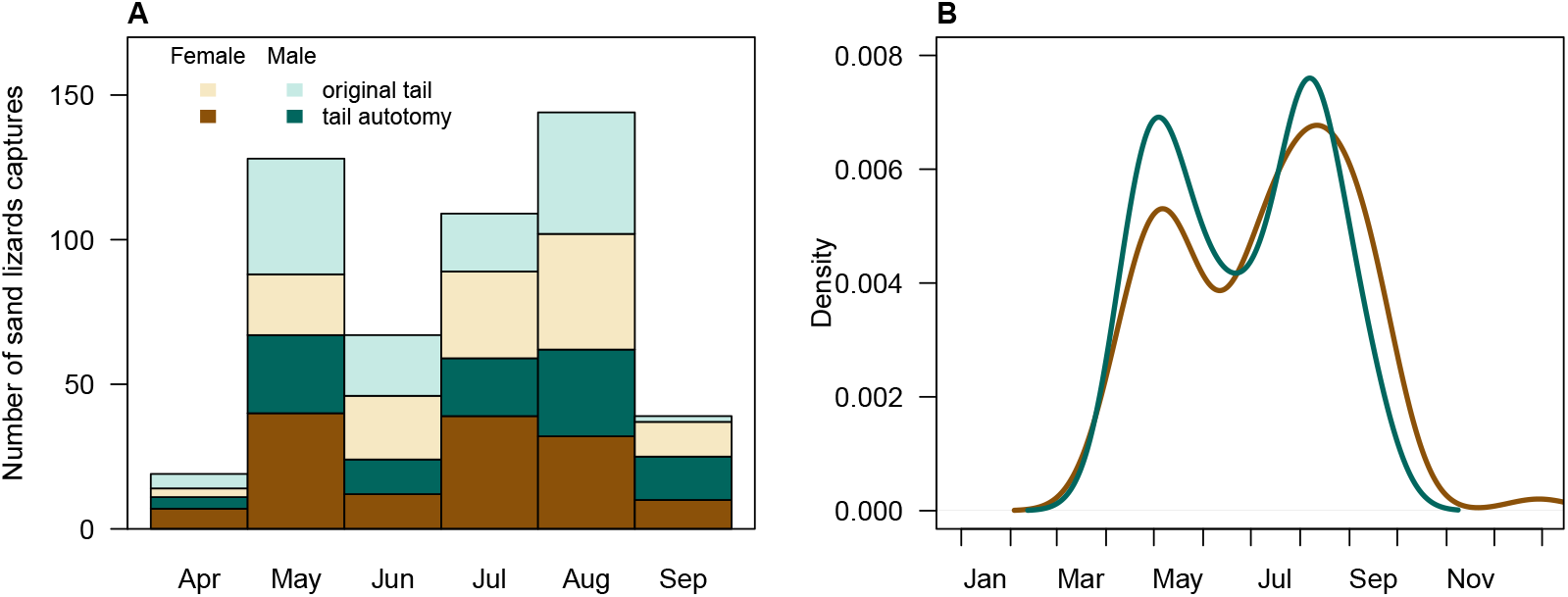
Seasonality of tail autotomy in sand lizards, *Lacerta agilis*. A. Observed number of captures of animals with their original tail and with a tail regenerate. B. Density of extrapolated dates when adult sand lizards experienced tail autotomy. The date of the tail autotomy was projected backward from the length of the tail regenerate on the day of capture, respecting the active season and hibernation periods.

The census sex ratio in young sand lizards was 0.500 (maximum likelihood confidence interval (CI) is [0.41, 0.59]; Fig. 3) and in adults, 0.453 (CI is [0.39, 0.51]), resulting in *O*_*sex*_ = 0.907 (Fisher’s exact test: *p* = 0.471). The frequency of tail autotomy in adult males was 0.627 (CI is [0.544, 0.71]) and in adult females 0.594 (CI is [0.52, 0.67]), giving *O*_*tail*_ = 1.055 (Fisher’s exact test: *p* = 0.609). In young animals, the frequency of tail autotomy in males was 0.167 (CI is [0.09, 0.28]) and in young females 0.333 (CI is [0.21, 0.48]), giving the odds of tail autotomy in young sand lizards as equal to 0.500 (Fisher’s exact test: *p* = 0.098).

**Figure 3:**
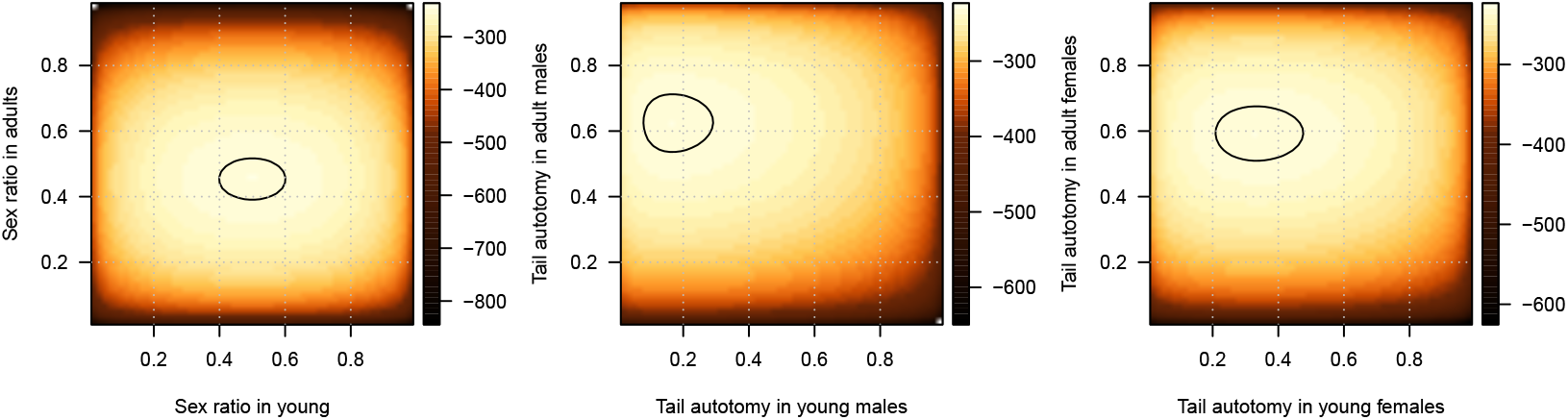
Marginal log-likelihood profiles of sex ratio and probability of tail autotomy in sand lizards. Shading indicates the log-likelihood of observing the data given the sex ratio (first pannel) or probability of tail autotomy (second and third pannel) indicated on the axes. Contours mark values up to 2 units lower than the maximum log-likelihood in the given test, indicating a confidence interval for the parameter value. Colour scale – log-likelihood.

The logistic regression model evaluating the increase in probability of tail autotomy with progressing approximate age and sex showed significant influence of the approximate age (Table 3, Fig. 4). The effect size of probability of tail autotomy increase with age is equal to 5.9% per month. The additive influence of sex and the interaction of the approximate age and sex did not significantly contribute to the model fit (Table 3).

**Table 3:** Logistic regression model evaluating change in the probability of tail autotomy with progressing age (in months) in sand lizard sexes (*Lacerta agilis*).

| Parameter | Modality | Estimate | St. error | $z$ statistic | $p$ |
| --- | --- | --- | --- | --- | --- |
| Intercept |  | -1.030 | 0.400 | -2.577 | 0.010 |
| Age |  | 0.057 | 0.019 | 2.963 | 0.003 |
| Sex | Male | -1.084 | 0.632 | -1.715 | 0.086 |
| Sex : Age | Male | 0.051 | 0.031 | 1.667 | 0.095 |

**Figure 4:**
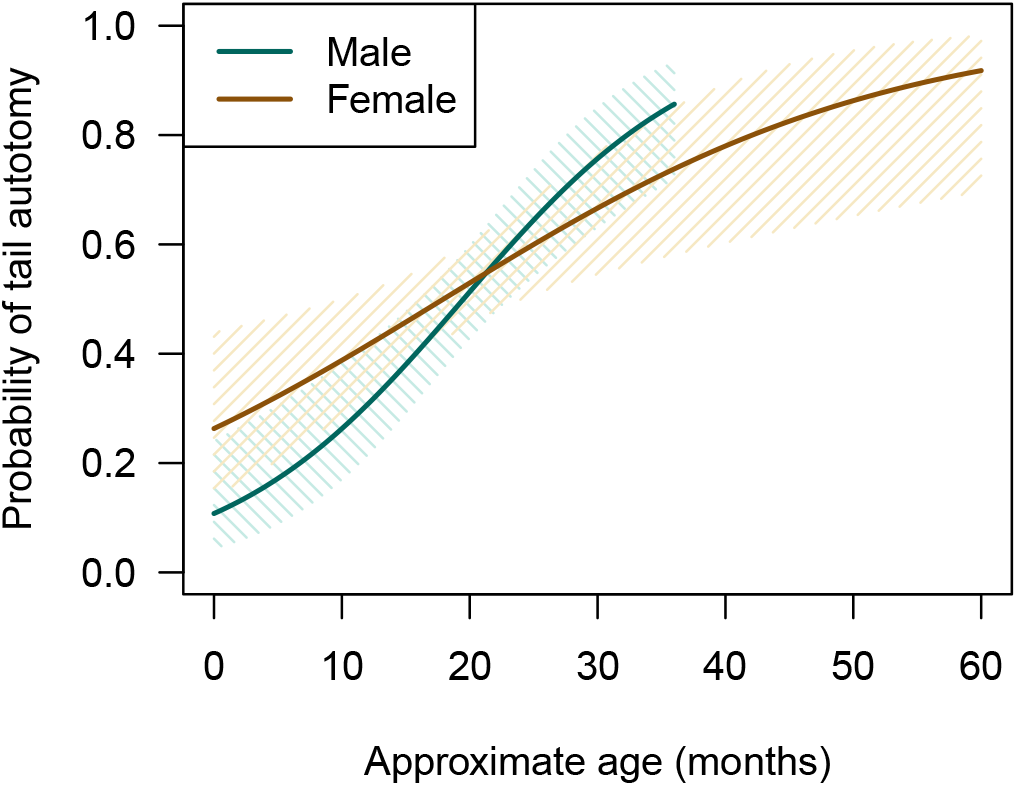
Partial fit of the logistic regression model evaluating the probability of tail autotomy with progressing approximate age in sand lizard sexes (*Lacerta agilis*). Solid lines – model prediction, shaded area – 95% prediction intervals.

The goodness-of-fit statistic for the CJS capture-recapture model was not significant (Test2: *χ*^2^ = 8.972, *df* = 68, *p >* 0.99; Test3: *χ*^2^ = 10.533, *df* = 48, *p >* 0.99, indicating that there is no apparent lack of fit in the data. The median *ĉ* was less then 1 (*ĉ* = 0.201, sampling SE = 0.669), and the survival rate and recapture probability parameters did not need to be corrected for overdispersion.

The CJS model that best fitted the observed capture-recapture data used survival rate dependent on animal sex and seasonal phases and a uniform recapture probability (Table S1). During the season phase I, female survival rate was significantly lower than in the season phase II (estimate ± SE: *ϕ*_phaseI_ = 0.009±0.009, *ϕ*_phaseII_ = 0.286±0.083; Fig. 5). In males, we observed a different pattern, where the survival rate was higher in season phase I than in phase II, but the difference was not significant in males (*ϕ*_phaseI_ = 0.086 ± 0.039, *ϕ*_phaseII_ = 0.006 ± 0.012; Fig. 5). The recapture probability remained statistically stable (*p*_*r*_ = 0.075 ± 0.007). The best model that considered sex as an explanatory variable without the seasonal element was significantly worse (ΔAIC_c_ = 19.206, *ϕ*_*F*_ = 0.103±0.036, *ϕ*_*M*_ = 0.069±0.030, *p*_*r*_ = 0.069 ± 0.007; Table S1).

**Figure 5:**
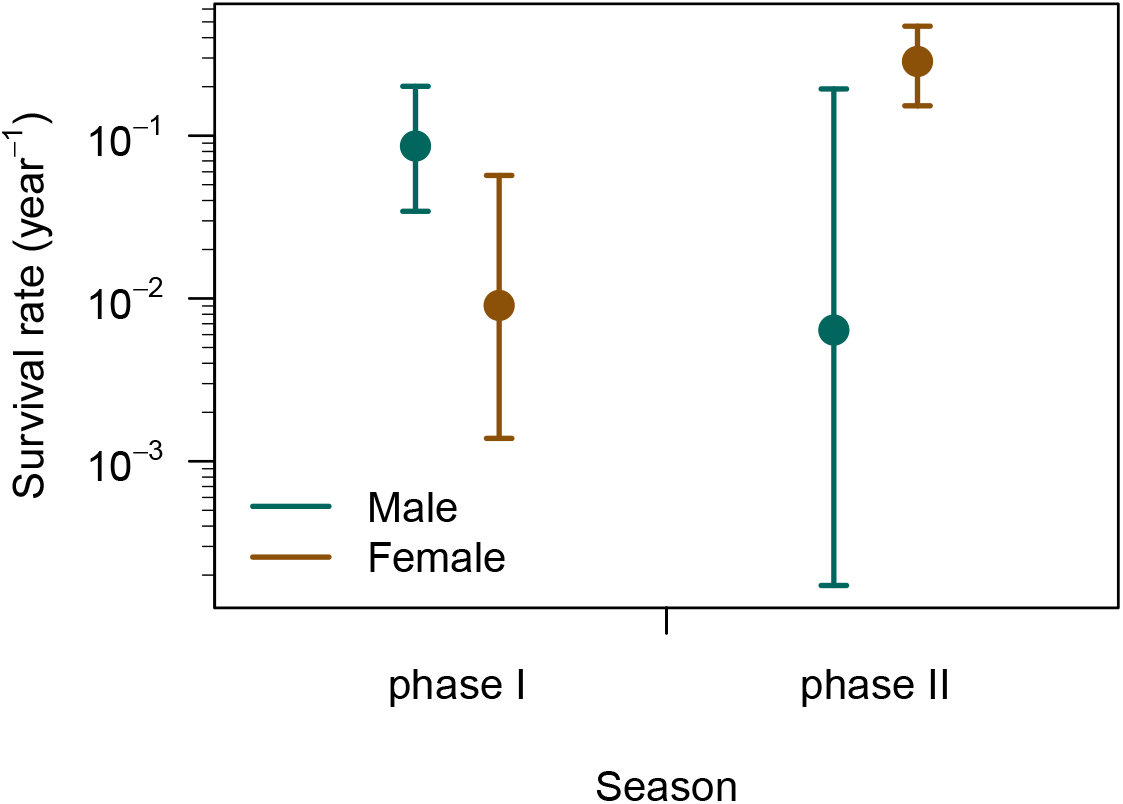
Survival rate of sand lizards (*Lacerta agilis*) based on the CJS capture-recapture model. The recapture probability was stable in the model and equal to 0.077 ± 0.008. Seasonal phase I represents the time characterised by mating behaviour and gestation, and phase II is the time with predominant resource accumulation prior to hibernation. Animals were categorized to seasonal phases based on the seasonal phase at the time of their first capture.

We recorded 180 tail autotomy events, where nine animals lost tails twice. To calculate the date of the tail autotomy event, we first estimated the seasonal growth rate of the tail regenerate from retraps of an individual with tail autotomy. The individual tail regenerate growth rate ranged from 0.016 to 1.417 mm day^*−*1^ during active season, and from 0.001 to 0.012 mm day^*−*1^ during hibernation (*n* = 27 and 2, respectively; Fig. S3). Projecting the tail regenerate length backwards, we calculated that most tail autotomy events occurred in August in males, and in May in females. Out of 179 tail autotomy events that were dated to the sand lizard active season, 86 occurred in season phase I (42 in males, 44 in females) and 93 in season phase II (40 in males, 53 in females). In adults (*n* = 144), the seasonal differences between sexes were not significant, although males had higher probability of a tail autotomy event in season phase I than females (odds ratio: 1.544, Fisher’s exact test: *p* = 0.242, Fig. 2B).

The set of equations estimating predation risk and escape rate (equations 4, 6, and 7) did not have an analytical solution for the observed values of the odds ratios and the survival rates. Using MCMC sampling with equation 8, we estimated that in sand lizards, predation risk is equal to 0.982 (highest posterior density interval (HPD) is [0.947, 0.999]) in males and 0.944 (HPD is [0.910, 0.972]) in females. The respective escape is 0.058 (HPD is [0.016, 0.071]) in males and 0.043 (HPD is [0.013, 0.074]) in females (Fig. 6). The MCMC runs converged according to the ln *L* trace and the potential scale reduction factors (PSRF) for all parameters (Gelman and Rubin 1992). The upper confidence limit of PSRF indicating covergence should be close to 1, and we observed values equal to 1.01 for all parameters.

**Figure 6:**
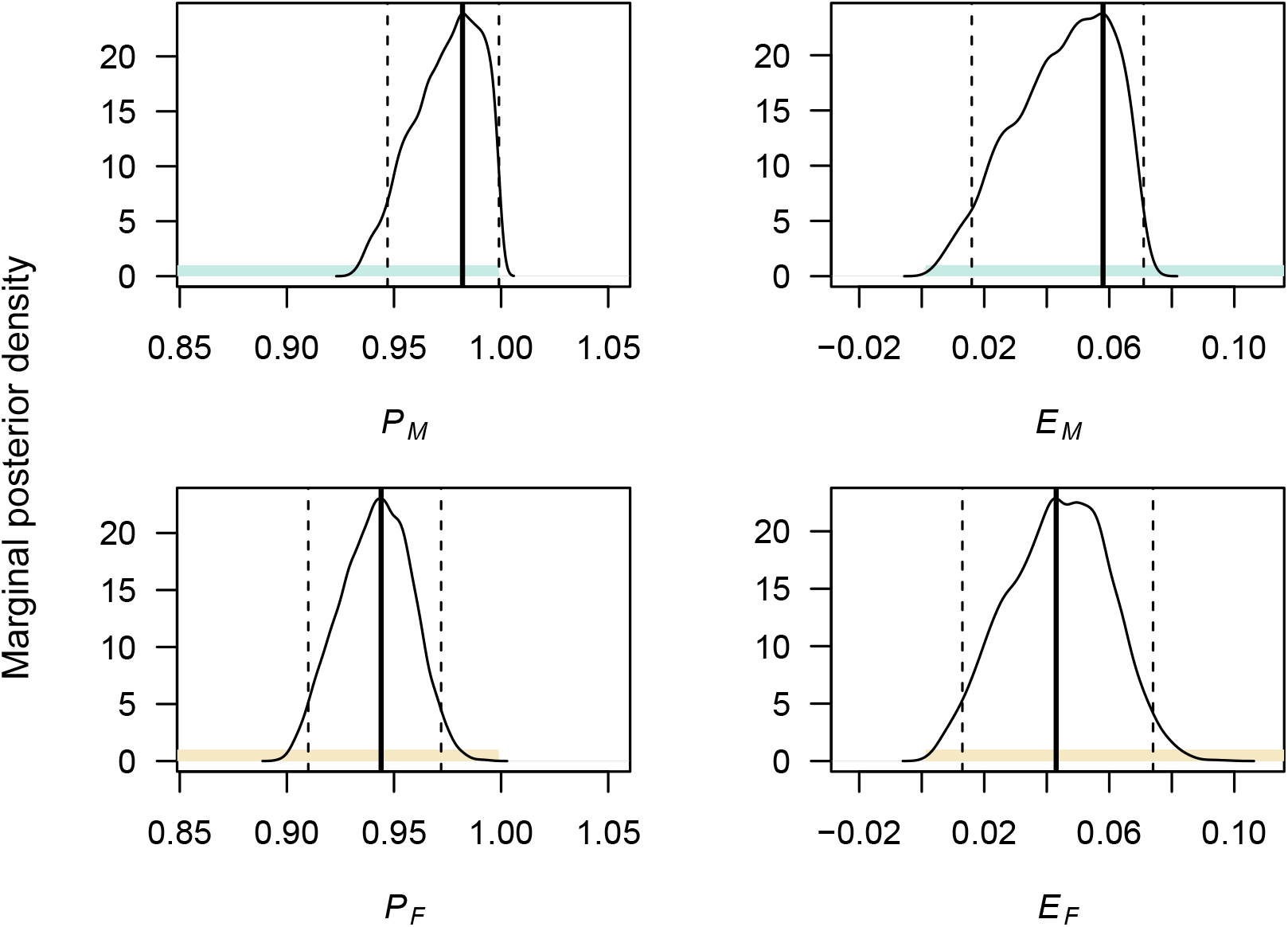
Marginal posterior density of predation risk (*P*) and escape rate (*E*) in sexes of sand lizards (*Lacerta agilis*). Coloured area – prior (green are males, brown females), dashed lines – highest posterior density interval at 95%, thick line – mode of the highest posterior density interval.

## Discussion

Theory suggests that visually-oriented predators will prey on conspicuous prey (Stevens and Merilaita 2011). In sand lizards, males in nuptial colouration signal their presence to attract mates and deter competitors (Olsson 1994; Bajer et al. 2011), and could thus face higher predation risk (Stuart-Fox et al. 2003; Marshall et al. 2015). We hypothesized that higher predation risk in males will manifest in changes in sex ratio, frequency of tail regenerates and survival depending on how successful males are at escaping the attack (Table 1). We observed adult sex ratio shifting in favour of females (Fig. 3), faster increase in probability of tail autotomy in males (Fig. 4), and sex-specific differences in seasonal survival (Fig. 5). Contrary to results in a grassland specialist lacertid *Takydromus viridipunctatus* (Lin et al. 2017) that perches on individual blades of grass (Chen et al. 2021), tail length is a poor predictor of sand lizard survival at our site (Table S1). Co-estimating sex-specific predation risk and escape rate showed that males tend to face a higher predation risk, but also have a higher escape rate (Fig. 6). However, none of our results were significant, and we conclude that predation risk and escape rate are statistically equal in sand lizard sexes.

The obvious reason for insignificant results is the lack of power in the test for the given effect size; the solution for this being an increase in sample size. For example, for Fisher’s exact test comparing young and adult sex ratio to be significant, we would have had to increase our sample size six times. This was not feasible in Hustopeče.

The result, where our data provided sufficient power, showed seasonal differences in female survival rate (Fig. 5). Accounting for recapture probability, we found a significantly lower survival rate in females first captured at the beginning of the active season (phase I), when sand lizards mate and lay eggs. Projection of the dates of tail autotomy events showed females experienced fewer attacks in season phase I than later in the active season (Fig. 2B). The odds of females experiencing a tail autotomy event in spring and early summer were 0.46 compared to males. Drivers of the low survival rate and low number of tail autotomy events in the first months after emergence from hibernation could be related to reproduction and environmental conditions during this time period.

We propose two non-exclusive mechanisms explaining the observed low survival and low number of tail autotomy events in season phase I in females. First, female locomotion is influenced by body allometry, and by increased resource requirements during gravidity (Olsson and Shine 1997; Olsson et al. 2000; Shine 2003; Kaliontzopoulou et al. 2013; Zamora-Camacho et al. 2014; Jagnandan and Higham 2018), making the females susceptible to predation. The relative clutch mass can exceed 50% of female weight in sand lizards (Roitberg et al. 2015). Females compensate for the increased load bearing through changes in biokinetics (Irschick et al. 2003) and modulation of thermal preferences (Le Galliard et al. 2003; López Juri et al. 2018), but our results show that at the site, the compensations are insufficient to support female survival during reproduction.

Second, low diurnal temperatures during spring and early summer months mean ectotherm lizards need more time to warm up (Amat et al. 2003). Sand lizards maintain body temperature range between 23 and 38^*°*^C (BÖhme and Bischoff 1984), but performance outside of their thermal optimum decreases their ability to escape. The trade-off between participating in reproduction and hiding to avoid predation risk seems to be skewed towards prioritizing reproduction, leading to low survival rate in season phase I in female sand lizards.

When developing our model (equations 4, 6 and 7), we considered only mortality attributable to predation (*m*_*P*_), setting mortality due to other causes at zero. While the solution reduced dimensionality of the problem, it deviated the model from reality. Mortality unrelated to predation contributes to the estimated survival rates, and could be sex- and season-dependent. Given the importance of seasonality in interaction with sex in modelling survival (Table S1), future developments of the predation risk and escape rate model (equation 8) should aim to include seasonal effects. In the current study, the projected dates of tail autotomy events (Fig. 2B) support the conclusion that predation contributes to low female survival in the sand lizard active season.

Tail autotomy can result not only from predation attempts but also from antagonistic intraspecific interactions. In fact, in island gecko populations, frequency of tail autotomy is negatively correlated with most predator indices (Itescu et al. 2017). Tail autotomy events associated with intraspecific aggression are more likely in males (reviewed in Bateman and Fleming 2009), and they could have increased frequency of tail autotomy events in males in season phase I and consequently increased the male escape rate in our results.

Reproduction is a costly endeavour (Shine 1980). The behavioural and morphological changes during reproduction attract attention of non-target receivers, the predators (Fowler-Finn and Hebets 2011). Our Bayesian modelling results show that tail autotomy events, likely attributable to an escape from a predator attack, improve survival of sand lizard sexes by about 5%. Tail autotomy events are most frequent in late spring and late summer, and males have a slightly higher predation risk. Gravid females do not lose their tail but their lives during a predator attack.

## Acknowledgements

We thank Adam Konečný for allowing us access to his orchard. This study was supported by the Institute of Vertebrate Biology of the Czech Academy of Sciences (RVO: 68081766), RECETOX Research Infrastructure (No. LM2018121) financed by the Ministry of Education, Youth and Sports, and the Operational Programme Research, Development and Education (the CETOCOEN EXCELLENCE project No. CZ.02.1.01/0.0/0.0/17_043/0009632).

## Authors’ Contributions

Radovan Smolinský, Zuzana Hiadlovská, and Natália Martínková conceptualised and designed the study; Radovan Smolinský, Zuzana Hiadlovská, Michal Škrobánek, and Natália Martínková collected data; Radovan Smolinský and Michal Škrobánek performed investigation; Zuzana Hiadlovská curated the data; Štěpán Maršala and Pavel Škrabánek designed and implemented the algorithm for sexing lizards; Natália Martínková performed statistical analyses; Radovan Smolinský, Pavel Škrabánek, and Natália Martínková wrote the manuscript, which all authors reviewed.

## Data Availability

The data is archived at the Dryad Data Repository (doi: 10.5061/dryad.q83bk3jm9, Smolinský et al. 2022).

## Supplementary material

Figure S1: Scheme of the ScalesCounter algorithm.

Figure S2: Sex determination in adult sand lizards, *Lacerta agilis*, using morphological isometry. A. Number of scales in the second row from the ventral medial line (ventralia) in sand lizards shows statistically significant sexual differences (*n* = 231). B. Receiver operating characteristic curve, indicating threshold number of ventralia scales that differentiate the sexes of sand lizards. At 58.5, the specificity of sex determination is equal to 0.85 and sensitivity to 0.86. The dashed line shows the ROC curve in young sand lizards whose sex was validated at recapture as adults (*n* = 59). C. Logistic regression model indicating probability of sex determination of sand lizards based on the number of ventralia scales. At 58.4, *p* = 0.5.

Figure S3: Individual seasonal growth rates of the tail regenerate in sand lizards (*Lacerta agilis*) conditional on tail regenerate length. winter – seasonal growth rate during hibernation

Table S1: Model selection of capture-recapture data of sand lizards (*Lacerta agilis*) analysed based on the CJS model. The models used group covariates sex (female, male), age (0young, 2adult), season (phase1, phase2), and individual covariates tailL (tail length corrected for body length) and aged (approximate age of the animal at first capture in months), Phi – survival rate, p – recapture probability, + – additive terms,: – interaction terms.

