## Supporting Information for "High predation risk decimates survival during the reproduction season"

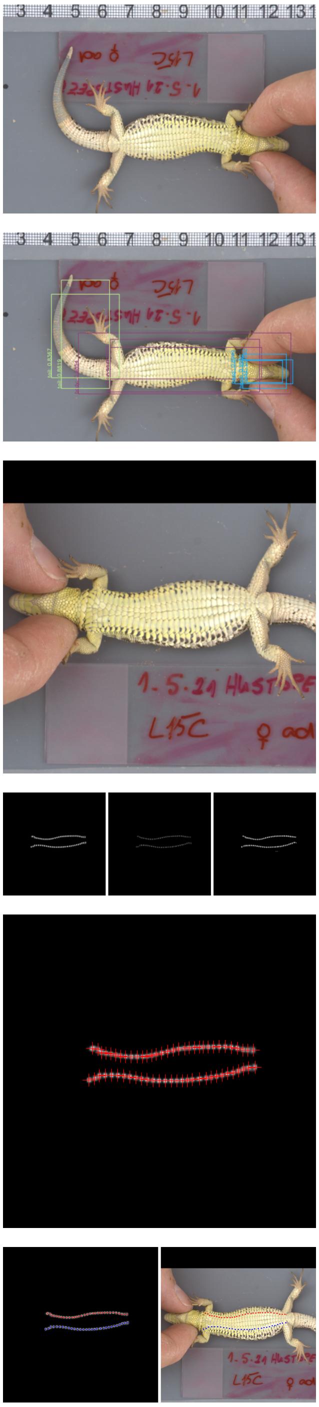

1. Photograph of sand lizard ventral side

2. Detection of lizard head, body and tail with the YOLO detector

3. Oriented and cropped photograph centered at the detected body; added zeros (black region) to allow a square crop

4. Three size- and shape-specific U-Net models detecting ventral scales in the second row from the medial line as circle gradients centered on the scale

5. Localization map of scale gradient centers, result of the first U-Net model that returned a valid answer (found 26–35 scales on each side)

6. Identification of the sides of the rows of scales based on their relative distance (left), and overlay of the detected scales localization map on the photograph (right)

Figure S1: Scheme of the ScalesCounter algorithm.

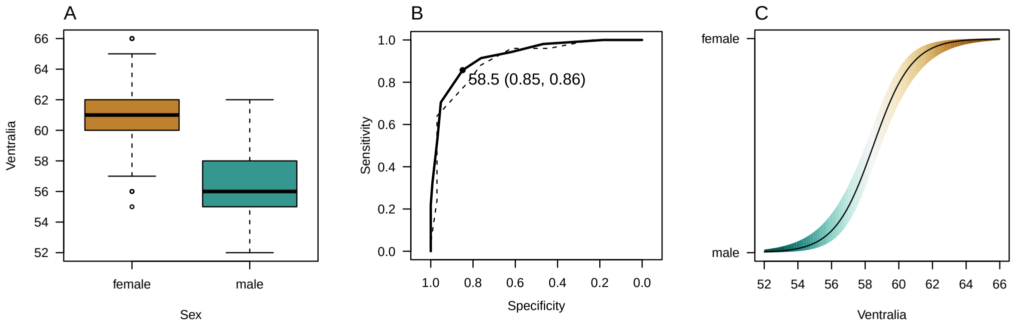


Figure S2: Sex determination in adult sand lizards, *Lacerta agilis*, using morphological isometry. A. Number of scales in the second row from the ventral medial line (ventralia) in sand lizards shows statistically significant sexual differences ($n = 231$). B. Receiver operating characteristic curve, indicating threshold number of ventralia scales that differentiate the sexes of sand lizards. At 58.5, the specificity of sex determination is equal to 0.85 and sensitivity to 0.86. The dashed line shows the ROC curve in young sand lizards whose sex was validated at recapture as adults ($n = 59$). C. Logistic regression model indicating probability of sex determination of sand lizards based on the number of ventralia scales. At 58.4, $p = 0.5$.

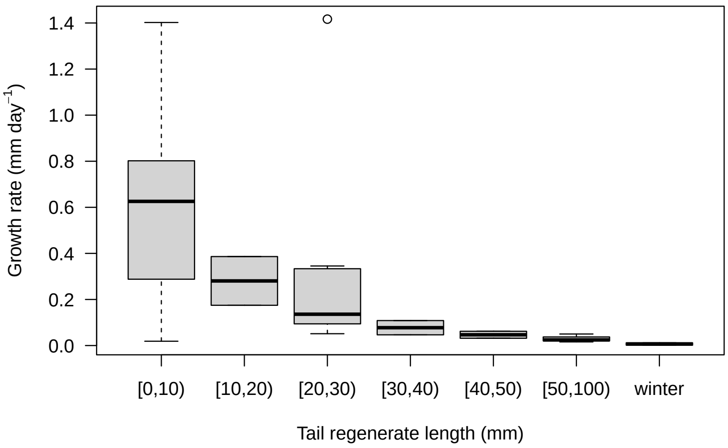


Figure S3: Individual seasonal growth rates of the tail regenerate in sand lizards (*Lacerta agilis*) conditional on tail regenerate length. winter – seasonal growth rate during hibernation.

Table S1: Model selection of capture-recapture data of sand lizards (*Lacerta agilis*) analysed based on the CJS model. The models used group covariates sex (female, male), age (0young, 2adult), season (phase1, phase2), and individual covariates tailL (tail length corrected for body length) and aged (approximate age of the animal at first capture in months), Phi – survival rate, p – recapture probability, + – additive terms, : – interaction terms.

| Model | $n_{par}$ | AICc | $\Delta$AICc | weight | Deviance |
|---|---|---|---|---|---|
| Phi(∼-1 +sex:season)p(∼1) | 5 | 1341.576 | 0.000000 | 3.462515e-01 | 971.2770 |
| Phi(∼-1 +sex:season)p(∼age) | 6 | 1342.979 | 1.402529 | 1.717261e-01 | 970.6303 |
| Phi(∼-1 +sex:season)p(∼sex) | 6 | 1343.393 | 1.817129 | 1.395748e-01 | 971.0448 |
| Phi(∼-1 +sex:season)p(∼season) | 6 | 1343.542 | 1.965829 | 1.295738e-01 | 971.1936 |
| Phi(∼-1 +sex:age:season)p(∼tailL) | 10 | 1345.318 | 3.742275 | 5.330487e-02 | 1324.8656 |
| Phi(∼-1 +sex:age:season)p(∼sex) | 10 | 1345.327 | 3.750975 | 5.307350e-02 | 964.6975 |
| Phi(∼-1 +sex:age:season)p(∼age) | 10 | 1345.841 | 4.264875 | 4.104741e-02 | 965.2114 |
| Phi(∼-1 +sex:age:season)p(∼season) | 10 | 1345.887 | 4.311375 | 4.010407e-02 | 965.2578 |
| Phi(∼-1 +sex:season)p(∼-1 +sex:season) | 8 | 1347.218 | 5.641982 | 2.061820e-02 | 970.7461 |
| Phi(∼-1 +sex:season)p(∼-1 +sex:age:season) | 12 | 1352.393 | 10.817029 | 1.550614e-03 | 967.5715 |
| Phi(∼-1 +sex:age:season)p(∼-1 +sex:age:season) | 16 | 1352.707 | 11.130534 | 1.325644e-03 | 959.3964 |
| Phi(∼season)p(∼1) | 3 | 1355.358 | 13.781982 | 3.521051e-04 | 989.1325 |
| Phi(∼season)p(∼sex) | 4 | 1356.888 | 15.311901 | 1.638531e-04 | 988.6298 |
| Phi(∼season)p(∼tailL) | 4 | 1357.057 | 15.480801 | 1.505838e-04 | 1348.9755 |
| Phi(∼season)p(∼age) | 4 | 1357.072 | 15.495901 | 1.494512e-04 | 988.8138 |
| Phi(∼season)p(∼season) | 4 | 1357.213 | 15.636501 | 1.393056e-04 | 988.9543 |
| Phi(∼aged)p(∼age) | 4 | 1357.637 | 16.061301 | 1.126482e-04 | 1349.5560 |
| Phi(∼aged)p(∼1) | 3 | 1357.693 | 16.116682 | 1.095718e-04 | 1351.6440 |
| Phi(∼season)p(∼-1 +sex:season) | 6 | 1357.975 | 16.399129 | 9.514063e-05 | 985.6269 |
| Phi(∼aged)p(∼tailL) | 4 | 1359.052 | 17.475701 | 5.553814e-05 | 1350.9704 |
| Phi(∼aged)p(∼sex) | 4 | 1359.234 | 17.657501 | 5.071238e-05 | 1351.1522 |
| Phi(∼1)p(∼1) | 2 | 1359.375 | 17.798692 | 4.725577e-05 | 995.1736 |
| Phi(∼aged)p(∼season) | 4 | 1359.527 | 17.950801 | 4.379502e-05 | 1351.4455 |
| Phi(∼age)p(∼1) | 3 | 1360.327 | 18.751082 | 2.935256e-05 | 994.1016 |
| Phi(∼sex)p(∼1) | 3 | 1360.782 | 19.205882 | 2.338231e-05 | 994.5563 |
| Phi(∼1)p(∼season) | 3 | 1360.873 | 19.296782 | 2.234337e-05 | 994.6473 |
| Phi(∼1)p(∼tailL) | 3 | 1360.918 | 19.341882 | 2.184517e-05 | 1354.8692 |
| Phi(∼aged)p(∼-1 +sex:season) | 6 | 1360.978 | 19.401529 | 2.120329e-05 | 1348.8061 |
| Phi(∼1)p(∼sex) | 3 | 1361.001 | 19.424582 | 2.096029e-05 | 994.7751 |
| Phi(∼tailL)p(∼1) | 3 | 1361.038 | 19.461582 | 2.057609e-05 | 1354.9889 |
| Phi(∼1)p(∼age) | 3 | 1361.301 | 19.725082 | 1.803618e-05 | 995.0756 |
| Phi(∼age)p(∼age) | 4 | 1361.336 | 19.759601 | 1.772755e-05 | 993.0774 |
| Phi(∼sex)p(∼sex) | 4 | 1361.422 | 19.845501 | 1.698227e-05 | 993.1633 |
| Phi(∼age)p(∼tailL) | 4 | 1361.746 | 20.170101 | 1.443809e-05 | 1353.6648 |
| Phi(∼age)p(∼sex) | 4 | 1361.932 | 20.355901 | 1.315721e-05 | 993.6738 |
| Phi(∼age)p(∼season) | 4 | 1361.982 | 20.405701 | 1.283364e-05 | 993.7236 |
| Phi(∼1)p(∼-1 +sex:season) | 5 | 1362.089 | 20.513500 | 1.216023e-05 | 991.7905 |
| Phi(∼sex)p(∼-1 +sex:season) | 6 | 1362.146 | 20.569829 | 1.182252e-05 | 989.7976 |
| Phi(∼sex)p(∼tailL) | 4 | 1362.274 | 20.698301 | 1.108696e-05 | 1354.1930 |
| Phi(∼sex)p(∼season) | 4 | 1362.312 | 20.736101 | 1.087939e-05 | 994.0540 |
| Phi(∼tailL)p(∼season) | 4 | 1362.587 | 21.010601 | 9.484131e-06 | 1354.5053 |
| Phi(∼tailL)p(∼sex) | 4 | 1362.707 | 21.130901 | 8.930479e-06 | 1354.6256 |
| Phi(∼sex)p(∼age) | 4 | 1362.709 | 21.133401 | 8.919323e-06 | 994.4512 |
| Phi(∼-1 +sex:age:season)p(∼1) | 9 | 1362.716 | 21.140210 | 8.889008e-06 | 984.1698 |
| Phi(∼tailL)p(∼tailL) | 4 | 1362.864 | 21.287901 | 8.256246e-06 | 1354.7826 |
| Phi(∼tailL)p(∼age) | 4 | 1363.000 | 21.423901 | 7.713484e-06 | 1354.9186 |
| Phi(∼season)p(∼-1 +sex:age:season) | 10 | 1363.287 | 21.711375 | 6.680768e-06 | 982.6579 |
| Phi(∼aged)p(∼-1 +sex:age:season) | 10 | 1363.314 | 21.738375 | 6.591184e-06 | 1342.8617 |
| Phi(∼age)p(∼-1 +sex:season) | 6 | 1363.336 | 21.759829 | 6.520859e-06 | 990.9875 |
| Phi(∼tailL)p(∼-1 +sex:season) | 6 | 1363.762 | 22.185629 | 5.270399e-06 | 1351.5902 |
| Phi(∼age)p(∼-1 +sex:age:season) | 10 | 1366.996 | 25.419675 | 1.046115e-06 | 986.3662 |
| Phi(∼sex)p(∼-1 +sex:age:season) | 10 | 1367.087 | 25.511075 | 9.993832e-07 | 986.4576 |
| Phi(∼1)p(∼-1 +sex:age:season) | 9 | 1367.117 | 25.540610 | 9.847332e-07 | 988.5702 |
| Phi(∼tailL)p(∼-1 +sex:age:season) | 10 | 1368.953 | 27.376675 | 3.932078e-07 | 1348.5000 |
| Phi(∼-1 +sex:age:season)p(∼-1 +sex:season) | 12 | 1393.259 | 51.682729 | 2.073145e-12 | 1008.4373 |
| Phi(∼-1 +sex:season)p(∼tailL) | 6 | 1419.548 | 77.971729 | 0.000000e+00 | 1407.3763 |
